# A new and accurate continuum description of moving fronts

**DOI:** 10.1101/101824

**Authors:** S T Johnston, R E Baker, M J Simpson

**Affiliations:** Mathematical Sciences, Queensland University of Technology, Brisbane, Australia; Mathematical Institute, University of Oxford, Oxford, United Kingdom

**Keywords:** Moving fronts, random walk, cell migration, Allee effect

## Abstract

Processes that involve moving fronts of populations are prevalent in ecology and cell biology. A common approach to describe these processes is a lattice-based random walk model, which can include mechanisms such as crowding, birth, death, movement and agent-agent adhesion. However, these models are generally analytically intractable and it is computationally expensive to perform sufficiently many realisations of the model to obtain an estimate of average behaviour that is not dominated by random fluctuations. To avoid these issues, both mean-field and corrected mean-field continuum descriptions of random walk models have been proposed. However, both continuum descriptions are inaccurate outside of limited parameter regimes, and corrected mean-field descriptions cannot be employed to describe moving fronts. Here we present an alternative description in terms of the dynamics of groups of contiguous occupied lattice sites and contiguous vacant lattice sites. Our description provides an accurate prediction of the average random walk behaviour in all parameter regimes. Critically, our description accurately predicts the persistence or extinction of the population in situations where previous continuum descriptions predict the opposite outcome. Furthermore, unlike traditional mean-field models, our approach provides information about the spatial clustering within the population and, subsequently, the moving front.

## Introduction

Moving fronts feature ubiquitously throughout biological and ecological processes [1, 2, 3, 4, 5, 6, 7, 8, 9, 10, 11, 12]. The introduction of non-native species can result in a catastrophic invasion wave if the introduced species out-competes native fauna [2, 13]. For example, the cane toad *bufo marinus* was introduced to north-eastern Australia in 1935, and has subsequently invaded much of northern Australia due to a lack of natural predation [9, 14]. Similarly, malignant tumours spread through the invasion of previously-healthy tissue, such as glioma cells moving throughout the brain to form glioblastoma [1, 3, 12, 15].

Lattice-based random walk models that include crowding, birth, death, movement and agent-agent adhesion are commonly used to describe processes that involve moving fronts [6, 16, 17, 18, 19, 20, 21]. For example, these random walk models have been used to interpret *in vitro* cell biology experiments by considering the position of the leading edge of the cell front or the cell density profile [6, 19, 21]. Illien *et al*. [18] consider random walks in the context of single-file diffusion models of active transport, that is, transport that requires energy due to an opposing force. However, the stochastic nature of random walks makes it problematic to efficiently examine the collective behaviour of a population, as a large number of realisations of the random walk must be performed to reduce the influence of stochastic fluctuations. Furthermore, it is difficult to determine meaningful population behaviour through analysis of the discrete process. There is, therefore, considerable interest in approaches that are both analytically tractable and avoid the computational expense of repeated simulations.

A common technique to analyse random walk processes is to consider a deterministic, continuum approximation of the discrete process [19, 20, 22, 23, 24, 25, 26, 27, 28, 29, 30]. The standard approximation, known as a mean-field (MF) approximation, results in a partial differential equation (PDE) [19, 20, 25]. The resulting PDE is amenable to analysis but only provides an accurate approximation of the discrete process in an extremely limited set of parameter regimes where spatial correlations are weak [22, 24, 31]. Hence, using these approximations to model moving fronts may provide an inaccurate estimate of the velocity of the moving front if the spatial correlations are important. This inaccuracy could have significant consequences if, for example, the front velocity is used to make decisions about implementing ecological control measures. To address the influence of spatial correlations, alternative approximation techniques have been proposed [22, 24, 28, 32, 33]. The corrected mean-field (CMF) approximation, which explicitly describes pairwise correlations, results in a system of ordinary differential equations (ODEs) that accurately approximate the discrete process for a wider range of parameter regimes, compared to the MF approximation [22, 24]. However, the CMF description is still invalid in parameter regimes where spatial correlations are sufficiently strong [28]. Furthermore, the CMF cannot be used to study moving fronts, as the governing equations cannot be evaluated anywhere that has zero agent density [22], such as areas where the population has yet to invade. The chain-and-gap approach (C&G), proposed by Johnston *et al*. [28], considers the dynamics of groups of contiguous occupied and vacant sites, and results in a system of ODEs that provide an accurate approximation of the discrete process in all parameter regimes. These groups are termed *chains* and *gaps* for the contiguous occupied and vacant sites, respectively. Additionally, the C&G description provides information about the spatial clustering and patchiness present in the system. There is considerable interest in determining the influence of local spatial structure on the persistence of a species [34, 35]. However, the C&G description has previously only been applied to discrete processes that are, on average, spatially uniform [28]. As such, the C&G description is not currently suitable for describing processes that contain moving fronts.

Here the C&G description presented by Johnston *et al*. [28] is extended to incorporate spatial variation so that the description can be applied to moving fronts. We interpret the discrete process in terms of chains and gaps, noting that the left-most site in each chain or gap can occur at any lattice site. The corresponding system of ODEs is derived and presented, and we demonstrate that the numerical solution of the ODE system provides an accurate approximation of the average discrete behaviour in all cases, even in parameter regimes where both the MF and CMF descriptions are inaccurate. This allows for the robust prediction of whether a population persists or becomes extinct, as well as reliable estimates of the velocity of the moving front. In addition, for the first time, the C&G description has been extended to include rates of birth, death and movement that are dependent on the length of the chain an agent belongs to. Furthermore, the C&G description includes explicit information about the spatial clustering present within a moving front.

## Random walk model

We consider a one-dimensional lattice-based random walk model where each lattice site may be occupied by, at most, one agent [36]. The lattice is interpreted as a combination of groups of contiguous occupied and vacant sites [28]. Agents on the lattice undergo birth, death and movement events. These events occur at rates *P*_*p*_^*n*^, *P*_*d*_^*n*^ and *P*_*m*_^*n*^ per unit time, respectively, where *n*∈[1, *N*] is the length of the chain an agent belongs to and *N* is the total number of sites. During a potential birth event, an agent randomly selects a nearest-neighbour site and attempts to place a daughter agent at that site. The birth event is successful provided that the target site is vacant [24]. During a death event, an agent is removed from the lattice [24]. During a potential movement event, an agent selects a nearest-neighbour site and attempts to move to that site, and is successful provided that the target site is vacant [24]. The target site selection is unbiased if both nearest-neighbour sites are vacant. If one nearest-neighbour site is occupied, the vacant nearest-neighbour site is selected with probability (1 − α)/2, where α ∈ [−1, 1] [26]. The constant parameter *α* represents the strength of agent-agent adhesion/repulsion. Setting *α* = 0 means that there is no agent-agent adhesion/repulsion, whereas setting *α*=*̸*0 simulates agent-agent adhesion (*α >*0) or repulsion (*α <*0) [26]. Note that *α* does not depend on the chain length.

Due to crowding, the success of birth and movement events depends on whether an agent has zero, one or two nearest-neighbour agents. These agents are referred to as *single*, *edge* or *middle* agents, respectively [28]. An example lattice configuration highlighting the different types of agents is presented in Figure 1. The necessary information to obtain the average numbers of single, edge and middle agents at each site *i* ∈ [1,*N*] is encoded within the average number of chains of length *n* ∈ [1,*N*] that contain the site *i*. The average number of single, edge and middle agents at site *i* are denoted *N*_*i*_^*S*^(*t*), *N*_*i*_^*E*^(*t*) and *N*_*i*_^*M*^(*t*), respectively. The average number of chains at time *t* where the left-most agent in the chain is at site *i*∈[1, *N*] is denoted *C*_*i*_^1^(*t*). Similarly, the average number of gaps at time *t*where the left-most vacant site in the gap is at site *i* ∈ [1, *N*] and the length is *m*∈[1, *N*-*i*+1]is denoted *G*^*m*^_*i*_(*t*). The spatially-dependent restriction on the length is due to the choice of no-flux boundary conditions. Note that *N*_*i*_^*S*^(*t*), *N*_*i*_^*E*^(*t*), *N*_*i*_^*M*^(*t*), *C*_*i*_^*n*^(*t*) and *G*^*m*^_*i*_(*t*) are all temporally-dependent but, for convenience, we do not explicitly include this dependence. The average numbers of edge, middle and single agents at site *i* are given by:

**Figure 1.**
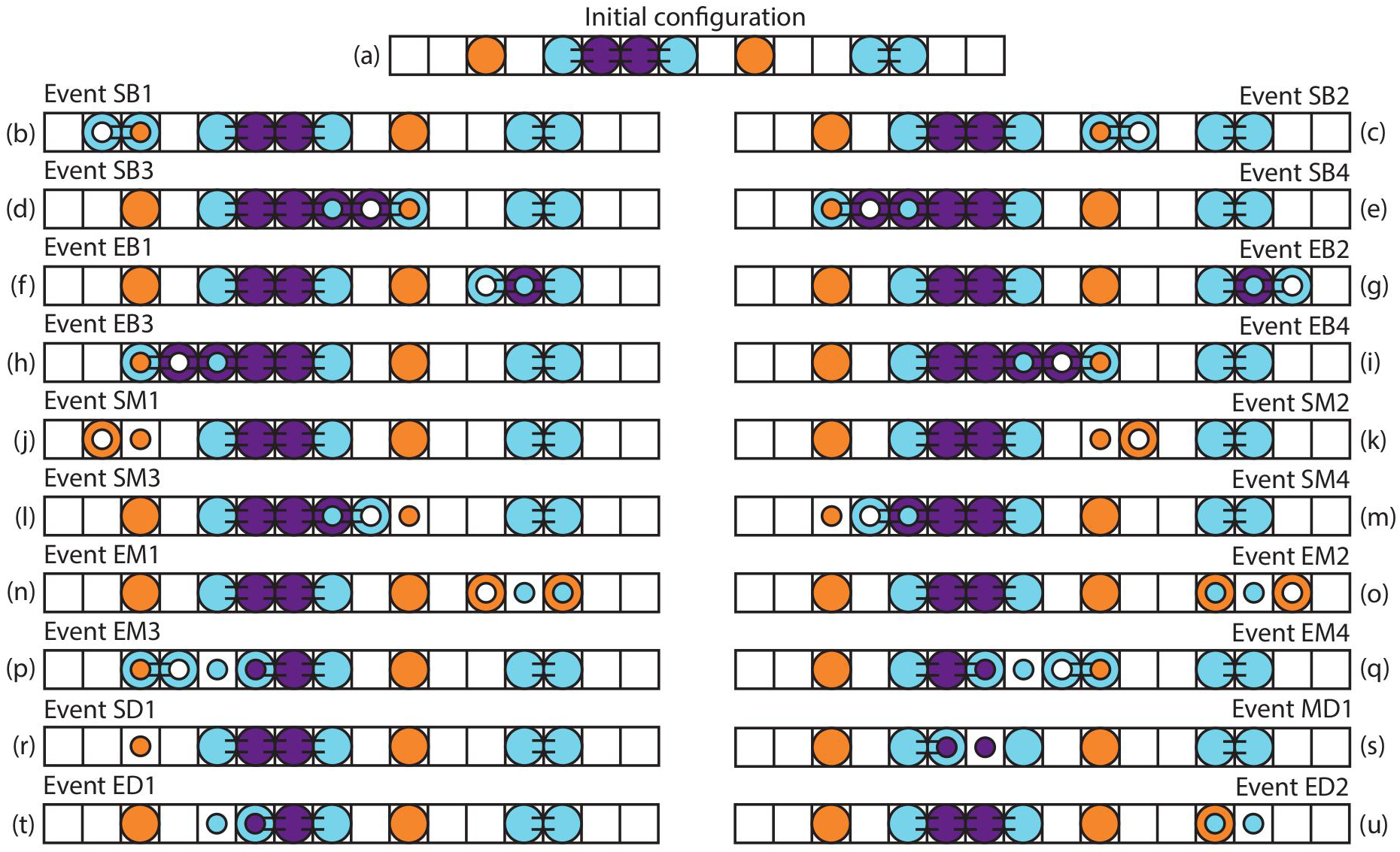
(a) Example lattice configuration with single (orange), edge (cyan) and middle (purple) agents. (b)-(u) Potential birth, death and movement events for the configuration of agents in (a), with subsequent change in agent type. Inset circles denote the agent type and location before the birth, death or movement event occurred. Lines connecting agents represent agent-agent adhesion/repulsion.

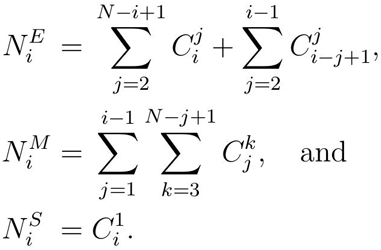

Note that middle agents cannot exist at sites *i* = 1 and *i* = *N* due to the choice of boundary conditions.

Similar to the approach of Johnston *et al.* [28], we consider how birth, death and movement events change the location and lengths of the chains and gaps, rather than the occupancy of an individual lattice site. This approach avoids making an assumption about the probability that a particular site is occupied or vacant, as the sites either side of a chain or gap are necessarily vacant or occupied, respectively. There are twenty different types of events that change either the location of the left-most agent in a chain, the length of a chain, or both. An example of each event is presented in Figure 1 for a particular configuration of agents. Events can have more than one potential outcome. For example, the potential outcomes of a single agent at site *i* undergoing a birth event can be classified into four groups. The daughter agent can be placed at site *i*−1 or *i*+1, and the gap that the daughter agent is placed in can be length one or greater. Here we detail each possible event and the subsequent change in configuration with respect to the number of chains and gaps. Events are referred to by a nomenclature describing the type of agent undergoing the event and the mechanism of the event itself, followed by a number highlighting the potential for multiple outcomes to arise from a specific event. For example, birth events for single agents are referred as SB events. Since there are four different types of SB events, we refer to these as SB1, SB2, SB3 and SB4 events. The details of each event are as follows:

**Event SB1:** A single agent at site *i* places a daughter agent at site *i* − 1, where the gap that includes site *i* − 1 is greater than length one. *C*_*i*_^1^ and *G*^*i-j*^_*j*_ decrease, *C*_*i*-1_^2^ and *G*_*j*_^*i-j*-1^ increase (Figure 1(b)).

**Event SB2:** A single agent at site *i* places a daughter agent at site *i* + 1, where the gap that includes site *i* + 1 is greater than length one. *C*_*i*_^1^ and *G*^*j*^_*i*+1_ decrease, *C*_*i*_^2^ and *G*_*i*+2_^*j*-1^ increase (Figure 1(c)).

**Event SB3:** A single agent at site *i* places a daughter agent at site *i*−1, where the gap that includes site *i*−1 is of length one. *C*_*i*_^1^, *G*^1^_*i*-1_ and *C*_*j*_^*i-j*-1^ decrease, *C*_*j*_^*i-j*+1^ increases (Figure 1(d)).

**Event SB4:** A single agent at site *i* places a daughter agent at site *i* + 1, where the gap that includes site *i*+1 is of length one. *C*_*i*_^1^, *G*^1^_*i*+1_ and *C*_*i*+2_^*j*^ decrease, *C*_*i*_^*j*+2^ increases (Figure 1(e)).

**Event EB1:** An edge agent at site *i* places a daughter agent at site *i* − 1, where the gap that includes site *i*−1 is greater than length one. *C*_*i*_^*j*^ and *G*^*i−k*^_*k*_ decrease, *C*_*i*−1_^*j*+1^ and *G*^*i*−1−*k*^_*k*_ increase, where *j* ≥ 2 (Figure 1(f)).

**Event EB2:** An edge agent at site *i* places a daughter agent at site *i* + 1, where the gap that includes site *i*+1 is greater than length one. *C*_*j*_^*i-j*+1^ and *G*^*k*^_*i*+1_ decrease, *C*_*j*_^*i-j*+2^ and *G*^*k*^_*i*+2_^1^ increase, where *j* ≥ 2 (Figure 1(g)).

**Event EB3:** An edge agent at site *i* places a daughter agent at site *i* − 1, where the gap that includes site *i*−1 is of length one. *C*_*i*_^*j*^, *G*^1^_*i*-1_ and *C*_*k*_^*i-k*-1^ decrease, *C*_*k*_^*i*+*j-k*^ increases, where *j* ≥ 2 (Figure 1(h)).

**Event EB4:** An edge agent at site *i* places a daughter agent at site *i* + 1, where the gap that includes site *i* + 1 is of length one. *C*_*j*_^*i-j*+1^, *G*^1^_*i*+1_ and *C*_*i*+2_^*k*^ decrease, *C*_*j*_^*i-j*+*k*+2^ increases, where *j* ≥ 2 (Figure 1(i)).

**Event SM1:** A single agent at site *i* moves to site *i* − 1, where the gap that includes site *i*−1 is greater than length one. *C*_*i*_^1^, *G*^*i-j*^_*j*_ and *G*^*k*^_*i*+1_ decrease, *C*_*i*-1_^1^, *G*^*i-j*-1^_*j*_ and *G*^*k*+1^_*i*_ increase (Figure 1(j)).

**Event SM2:** A single agent at site *i* moves to site *i* + 1, where the gap that includes site *i*+1 is greater than length one. *C*_*i*_^1^, *G*^*i-j*^_*j*_ and *G*^*k*^_*i*+1_ decrease, *C*_*i*+1_^1^, *G*_*j*_^*i-j*+1^ and *G*^*k*-1^_*i*+2_ increase (Figure 1(k)).

**Event SM3:** A single agent at site *i* moves to site *i* − 1, where the gap that includes site *i* − 1 is of length one. *C*_*i*_^1^, *G*^1^_*i*-1_, *G*^*j*^_*i*+1_ and *C*_*k*_^*i-k*-1^ decrease, *C*_*k*_^*i-k*^ and *G*^*j*+1^_*i*_ increase (Figure 1(l)).

**Event SM4:** A single agent at site *i* moves to site *i* + 1, where the gap that includes site *i*+1 is of length one. *C*_*i*_^1^, *G*^1^_*i*+1_, *G*^*i-j*^_*j*_ and *C*_*i*+2_^*k*^ decrease, *C*_*i*+1_^*k*+1^ and *G*^*i-j*+1^_*j*_ increase (Figure 1(m)).

**Event EM1:** An edge agent at site *i* moves to site *i* − 1, where the gap that includes site *i*−1 is greater than length one. *C*_*i*_^*j*^ and *G*^*i-k*^_*k*_ decrease, *C*_*i*-1_^1^, *G*^1^_*i*_, *C*_*i*+1_^*j*-1^ and *G*_*k*_^*i-k*-1^ increase, where *j* ≥ 2 (Figure 1(n)).

**Event EM2:** An edge agent at site *i* moves to site *i* + 1, where the gap that includes site *i* + 1 is greater than length one. *C*_*j*_^*i-j* + 1^ and *G*^*k*^_*i* + 1_ decrease, *C*_*i* + 1_^1^, *G*^1^_*i*_, *C*_*j*_^*i-j*^ and *G*^*k*-1^_*i*+2_ increase, where *j* ≥ 2 (Figure 1(o)).

**Event EM3:** An edge agent at site *i* moves to site *i* − 1, where the gap that includes site *i*−1 is of length one. *C*_*i*_^*j*^, *G*^1^_*i* − 1_ and *C*_*k*_^*i−k*−1^ decrease, *C*_*i*+1_^*j*-1^, *G*^1^_*i*_ and *C*_*k*_^*i−k*^ increase, where *j* ≥ 2 (Figure 1(p)).

**Event EM4:** An edge agent at site *i* moves to site *i* + 1, where the gap that includes site *i*+1 is of length one. *C*_*j*_^*i−j*+1^, *G*^1^_*i*+1_ and *C*_*i*+2_^*k*^ decrease, *C*_*j*_^*i−j*^, *G*^1^_*i*_ and *C*_*i*+1_^*k*+1^ increase, where *j* ≥ 2 (Figure 1(q)).

**Event SD1:** A single agent at site *i* dies. *C*_*i*_^1^, *G*^*i−j*^_*j*_ and *G*^*k*^_*i*+1_ decrease, *G*_*j*_^*i*+*k−j*+1^ increases (Figure 1(r)).

**Event MD1:** A middle agent at site *i* dies. *C*_*j*_^*k*^ decreases, *C*_*j*_^*i−j*^, *C*_*i*+1_^*j*+*k−i*-1^ and *G*^1^_*i*_ increase, where *j* ≥ 3 (Figure 1(s)).

**Event ED1:** An edge agent at site *i* dies, where site *i* + 1 is occupied and site *i* − 1 is vacant. *C*_*i*_^*j*^ and *G*^*i-k*^_*k*_ decrease, *C*_*i*+1_^*j*-1^ and *G*_*k*_^*i-k*+1^ increase, where *j* ≥ 2 (Figure 1(t)).

**Event ED2:** An edge agent at site *i* dies, where site *i* − 1 is occupied and site *i* + 1 is vacant. *C*_*j*_^*i-j*+1^ and *G*^*k*^_*i*+1_ decrease, *C*_*j*_^*i−j*^ and *G*^*k*+1^_*i*_ increase, where *j* ≥ 2 (Figure 1(u)).

To obtain expressions for the time rate of change of *C*_*i*_^*n*^ and *G*^*m*^_*i*_, we consider the rate at which each event occurs at site *i*, and all possible results of each event. Birth events are always successful for single agents and, as such, occur at rate *P*_*p*_^1^*C*_*i*_^1^. Similarly, movement events for single agents are always successful and are not influenced by agent-agent adhesion/repulsion. Therefore single agent movement events occur at rate *P*_*m*_^1^*C*_*i*_^1^. Birth events for edge agents are, on average, unsuccessful half the time due to crowding. Hence, birth events occur at rates 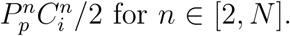 Movement events for edge agents are influenced by both crowding and agent-agent adhesion/repulsion, and occur at rates 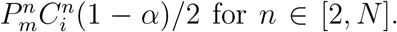 Neither birth or movement events can occur for middle agents. For all agent types, death events are not influenced by crowding and occur at rates 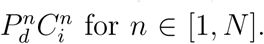 Note that all rates are chain-length dependent. Practical examples that could be described using chain-length dependent rates arise in a variety of situations [28, 37]. For example, Hedayati *et al.* [37] demonstrate that nanoparticle-mediated heating causes cytotoxicity in prostate cancer cells, provided the volume of cells is above a threshold value. Furthermore, the cytotoxicity increases with the cell number. Our framework would be suitable for modelling this process, as we are able to impose death rates that are zero below a threshold length, and an increasing function with respect to length otherwise.

While the rates at which events occur for each mechanism and agent type combination are known, multiple events can occur for a mechanism and agent type combination. For example, there are four types of birth events for single agents and, as such, we require the proportion of single birth events that are SB1, SB2, SB3 and SB4 events. For a single birth event to be an SB1 event, the agent at site *i* must place a daughter agent at site *i* − 1 and the gap that contains site *i* − 1 must be of length two or greater. Note that the gap cannot include site *i*, as the selected agent occupies that site. The proportion of single birth events where the agent at site *i* selects a target site at *i* − 1 is 1/2. The proportion of single birth events where the agent selects a target site that is part of a gap of length two or greater is 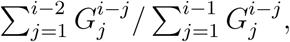 that is, the number of gaps including site *i* − 1 that are of length two or greater divided by the total number of gaps including site *i* − 1. Hence the rate at which SB1 events occur at site *i*, and subsequently decrease *C*^1^_*i*_ and increase *C*_*i*−1_^2^, is 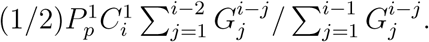 SB1 events also decrease *G*_*j*_^*i-j*^ and increase *G*_*j*_^*i-j*-1^, where *j*∈[1, *i*-2] The proportion of SB1 events that change *G*_*j*_^*i-j*^ and *G*_*j*_^*i-j*-1^ for a specific *j*∈[1, *i*-2] is 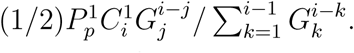 Therefore, we can determine the expected rate of change for all chains and gaps affected by an SB1 event at site *i*. Following a similar process for all events we obtain transition rates for *C*_*i*_^*n*^ and *G*^*m*^_*i*_, where *i* ∈ [1,*N*], *n* ∈ [1,*N* − *i* + 1] and *m* ∈ [1,*N* − *i* + 1]. The resulting system of ODEs is presented in Appendix A.1 for *C*_*i*_^*n*^ and Appendix A.2 for *G*^*m*^_*i*_.

## Traditional mean-field descriptions

Traditional MF descriptions of lattice-based random walk models containing crowding, birth, death, movement and agent-agent adhesion do not have the flexibility to describe processes where the rate of birth, death and/or movement is arbitrarily chain-length dependent. However, with certain simplifying assumptions, MF descriptions of the discrete process can be derived [20, 38]. Here we examine two special cases of the discrete process, where continuum MF descriptions have been presented previously.

Special case 1: One simplifying assumption is that the birth, death and movement rates are independent of chain length. Hence we define *P*_*p*_ = *P*_*p*_^1^ = *P*_*p*_^2^ = *…* = *P*_*p*_^*N*^, *P*_*d*_ = *P*_*d*_^1^ = *P*_*d*_^2^ = *…*= *P*_*d*_^*N*^, and *P*_*m*_ = *P*_*m*_^1^ = *P*_*m*_^2^ = *…*= *P*_*m*_^*N*^. There is, therefore, no positive or negative benefit associated with an agent being near other agents. Furthermore, there is no agent-agent adhesion/repulsion, and hence *α* = 0. The MF description for this case takes the form of a reaction-diffusion equation, known as the Fisher-Kolmogorov model [20, 38, 39, 40],

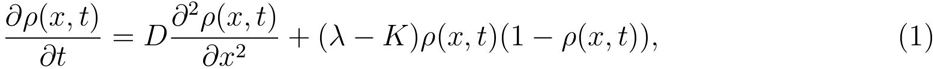

where

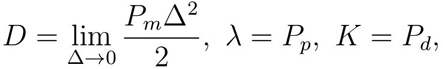

and ρ(*x*, *t*)is the agent density [20, 38].

Special case 2: An alternative simplifying assumption is that birth, death and movement rates depend on whether an agent is isolated or not. An isolated agent has zero nearest-neighbours [38], corresponding to a chain of length one. All grouped agents, that is, agents with at least one nearest-neighbour, undergo birth, death and movement events at the same rate. Grouped agents correspond to agents that are part of a chain of length two or greater. Hence we define *P*_*p*_^*G*^ = *P*_*p*_^2^ = *…* = *P*_*p*_^*N*^, *P*_*d*_^*G*^ = *P*_*d*_^2^ = *…*= *P*_*d*_^*N*^, and *P*_*m*_^*G*^ = *P*_*m*_^2^ = *…*= *P*_*m*_^*N*^. This assumption introduces a potential positive or negative benefit associated with an agent being adjacent to other agents. For example, if *P*_*d*_^1^*> P*_*d*_^*G*^ then isolated agents are more likely to die, compared to other agents, and hence there is a positive benefit associated with being adjacent to other agents. This allows for significant flexibility in modelling a variety of competitive or co-operative processes [38]. Again, there is no agent-agent adhesion/repulsion, and α = 0. The MF description for this case is a reaction-diffusion equation with a nonlinear diffusivity function and Allee kinetics [38],

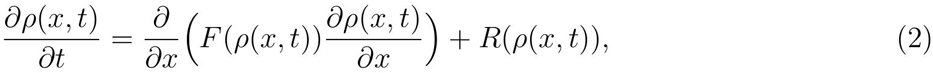

where

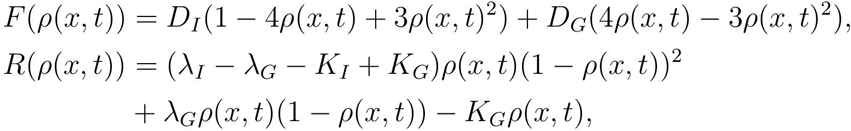

and

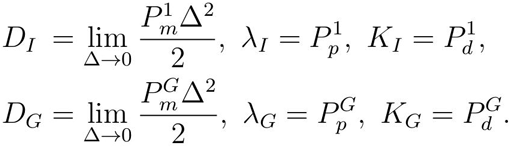

While CMF descriptions have been presented for certain lattice-based random walk models, these descriptions are unsuitable for studying problems containing moving fronts as the system of governing ODEs is singular at zero agent density [22, 24, 26].

## Results

The solution of the MF model leads to a prediction of the average agent density profile as a function of position and time, whereas the C&G description provides the number of chains and gaps of all possible lengths as a function of location and time. Hence to compare the MF descriptions with the C&G description, it is necessary to reconstruct the agent density at each location, *ρ*_*i*_, from *C*_*i*_^*n*^ and *G*^*m*^_*i*_. The agent density at site *i* is the sum of all possible chains that include site *i*, namely,

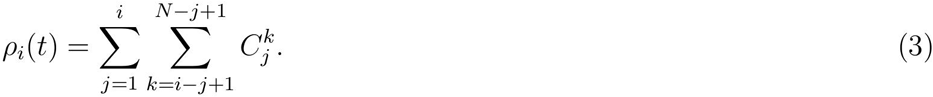

An example of the output from the one-dimensional discrete model, illustrating twenty identically-prepared realisations, is presented in Figures 2(a)-(c) at *t* = 0, *t* = 100 and *t*= 200, respectively. Identically-prepared realisations refer to simulations of the discrete model performed with the same initial condition, parameter regime and boundary conditions. Initially, the domain is fully-occupied for 61 ≤ *x* ≤ 140, and vacant otherwise. As time increases, the population spreads into the initially-vacant region. Note that Figures 2(a)-(c) each show twenty one-dimensional simulations, rather than one two-dimensional simulation. To obtain the average behaviour of the agent population, *M* identically-prepared realisations of the discrete model are performed and the binary lattice occupancy at each site *i*, ρˆ_*i*_, is calculated for each realisation. Note that the discrete model is simulated with the Gillespie algorithm [41]. The binary lattice occupancy is then averaged, giving 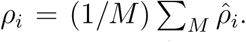The averaged density profile from the discrete model is presented in Figure 2(d), with the numerical solutions to both Equation (1) and the C&G governing equations superimposed, at *t* = 100 and *t* = 200. Both continuum descriptions match the averaged discrete model predictions extremely well. Details of the numerical techniques used to solve the PDEs and systems of ODEs are presented in Appendix B.

**Figure 2.**
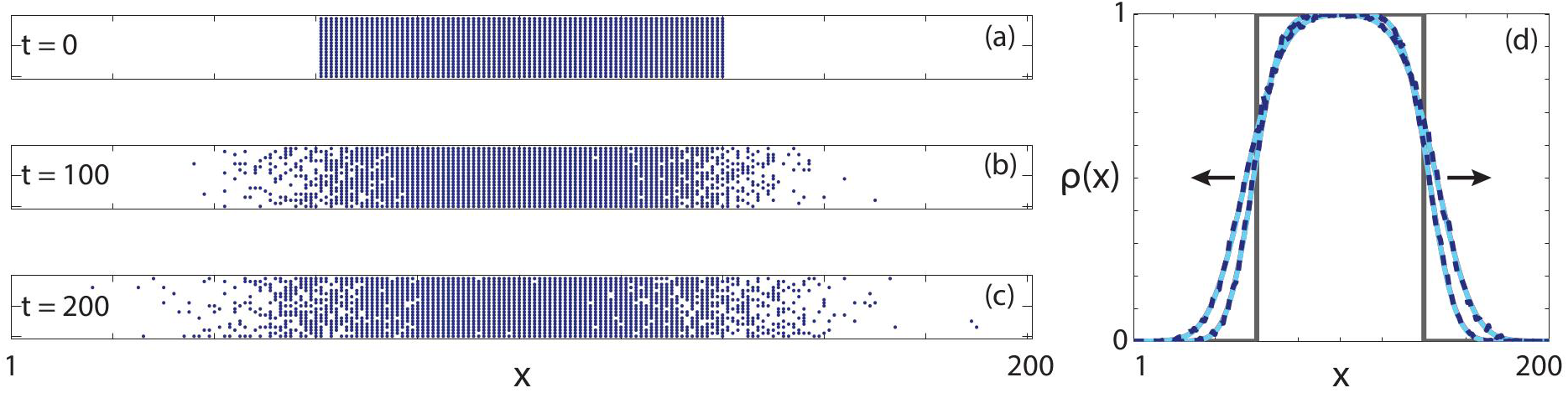
(a)-(c) Twenty identically-prepared realisations of the one-dimensional discrete model at (a) *t* = 0, (b) *t* = 100, and (c) *t* = 200, as indicated. (d) Comparisons of density profiles obtained from the averaged discrete model (blue, dashed), MF description (red, solid) and C&G description (cyan, solid). Note that the cyan and red curves are visually indistinguishable at this scale. All results are obtained using 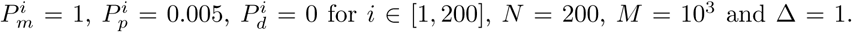. These parameter choices correspond to Special case 1. Profiles are presented at *t* = 100 and *t*= 200, and the arrow indicates the direction of increasing time. Grey lines representthe initial condition.

The parameter regime considered in Figure 2 has *P*_*m*_/*P*_*p*_ ≫ 1 and *P*_*d*_ = 0, and, as such, we expect the MF description to approximate the average discrete behaviour well since this combination of parameters avoids the formation of significant agent clustering [20, 22]. We now consider a parameter regime where 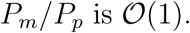 In such parameter regimes, the spatial correlations are significant and, subsequently, the MF description does not provide a valid approximation of the average discrete behaviour [22, 28]. To highlight this, a comparison between the average discrete behaviour and the numerical solution of Equation (1) is presented in Figure 3(a). For these results, the domain is initially fully-occupied for *x* ≤ 20, and is vacant otherwise. The MF description predicts a front that is significantly sharper, and has a higher front speed, compared to the discrete model. In contrast, the numerical solution of the C&G description predicts the average discrete behaviour well, matching both the shape and position of the averaged discrete data. Furthermore, the distribution of chain and gap lengths, *C*^*n*^ and *G*^*m*^, matches the observed average distribution of chains and gaps in the discrete model, as shown in Figures 3(b)-(c). As such, the C&G description provides an accurate estimate of the front shape and speed, as well as a valid prediction of the clustering of occupied and vacant sites in the system.

**Figure 3.**
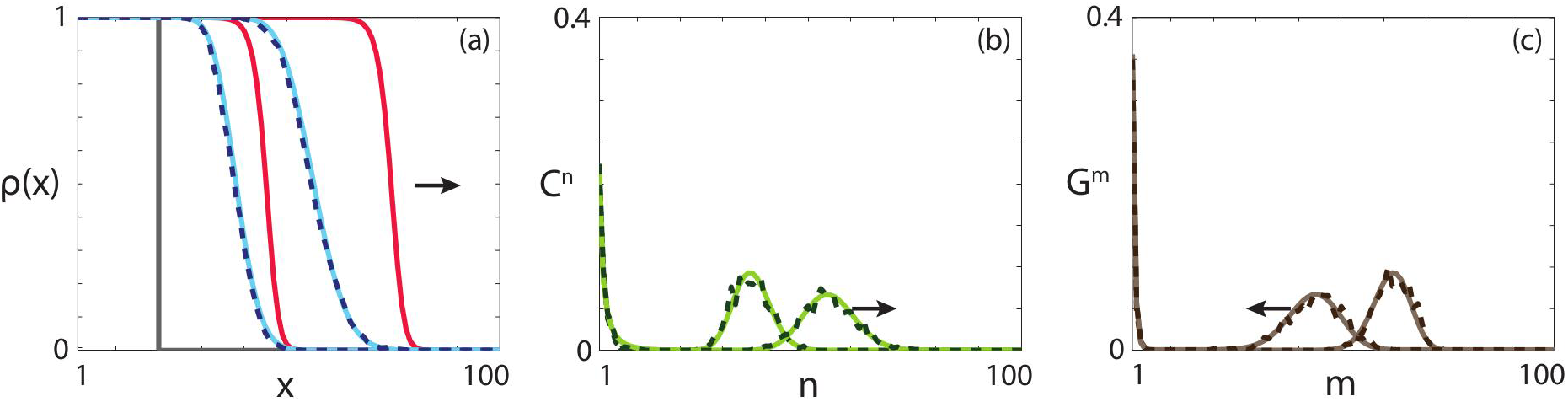
(a) Comparisons of density profiles obtained from the averaged discrete model (blue, dashed), MF description (red, solid) and C&G description (cyan, solid). (b) Comparison of the chain distribution obtained from the averaged discrete model (dark green, dashed) and C&G description (light green, solid). (c) Comparison of the gap distribution obtained from the averaged discrete model (dark brown, dashed) and C&G description (light brown, solid). All results are obtained using *P*_*m*_^*i*^ = 0.5 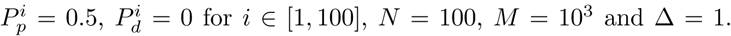. These parameter choices correspond to Special case 1. Profiles are presented at *t* = 100 and *t* = 200, and the arrow indicates the direction of increasing time. Grey lines represent the initial condition.

If the birth, death and movement rates depend on whether an agent has zero or at least one nearest-neighbour agent then the MF description of the discrete model is Equation (2) [38]. A comparison of the average discrete behaviour, the numerical solution of Equation (2) and the numerical solution of the C&G description in an appropriate parameter regime is presented in Figure 4(a). Interestingly, the MF description predicts that the agent population moves in the negative *x* direction, and would subsequently become extinct. In contrast, both the discrete model and the C&G description suggest that the agent population persists, and spreads in the positive *x* direction. Again, the C&G description matches the average discrete behaviour extremely well. The results in Figure 4(a) highlight the need for an accurate approximation. If the naive approach of implementing a MF approximation to describe the spread of an invasive species is taken, it might be recommended that no culling measures are required to curtail the spread of the species. Obviously, such a recommendation is incorrect if the aim is to halt the invasion of the alien species. The clustering present in the system is highlighted in Figures 4(b)-(c), for chains and gaps, respectively. Intuitively, as time increases and the population spreads, the average chain length increases and the average gap length decreases. The C&G description predicts both the average chain and gap distributions in the discrete model well.

**Figure 4.**
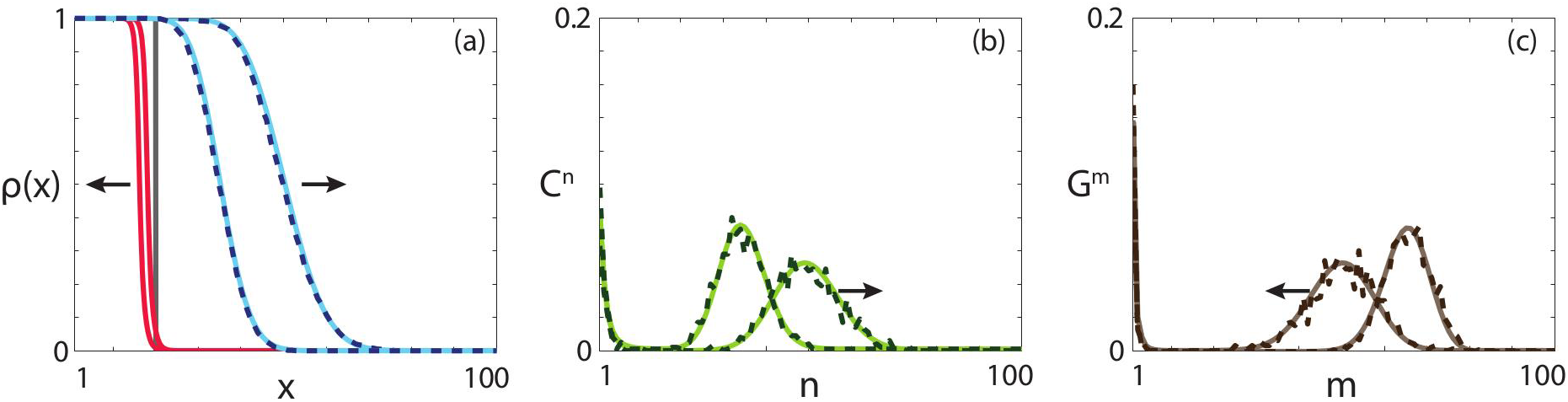
(a) Comparisons of density profiles obtained from the averaged discrete model (blue, dashed), MF description (red, solid) and C&G description (cyan, solid). (b)Comparison of the chain distribution obtained from the averaged discrete model (dark green, dashed) and C&G description (light green, solid). (c) Comparison of the gap distribution obtained from the averaged discrete model (dark brown, dashed) and C&G description (light brown, solid). All results are obtained using *P*_*m*_^1^ = 0.5, *P*_*p*_^1^= 0.4, *P*_*d*_^1^= 0.7, *P*_*m*_^*i*^= 0.25, *P*_*p*_^*i*^=0.3, *P*_*d*_^*i*^= 0 for *i*∈[2, 100]*N*=100, *M*= 10^3^and ∆ = 1. These parameter choices correspond to Special case 2. Profiles are presented at *t*=50 and *t*=100, and the arrow indicates the direction of increasing time. Grey lines represent the initial condition.

Introducing a non-zero death rate for chains of length two or greater reduces the carrying capacity in the MF description of the discrete model [38]. To determine whether this reduction is an accurate reflection of the average discrete behaviour, we present a comparison of the average discrete behaviour, and the numerical solutions of both Equation (2) and the C&G governing equations in Figure 5(a). Both the discrete model and the C&G description predict that the peak agent density near *x* = 50 decreases between *t* = 25 and *t* = 50, whereas the MF description predicts that the peak agent density at this location is approximately constant, at *ρ* = 0.646 [38], after *t* = 25. Interestingly, both the discrete model and the C&G description predict that the population eventually goes extinct. In contrast, the MF description predicts that the population persists and spreads throughout the domain. Again, these results highlight the importance of implementing an accurate approximation to obtain meaningful conclusions, as well as the robust nature of the C&G description.

**Figure 5.**
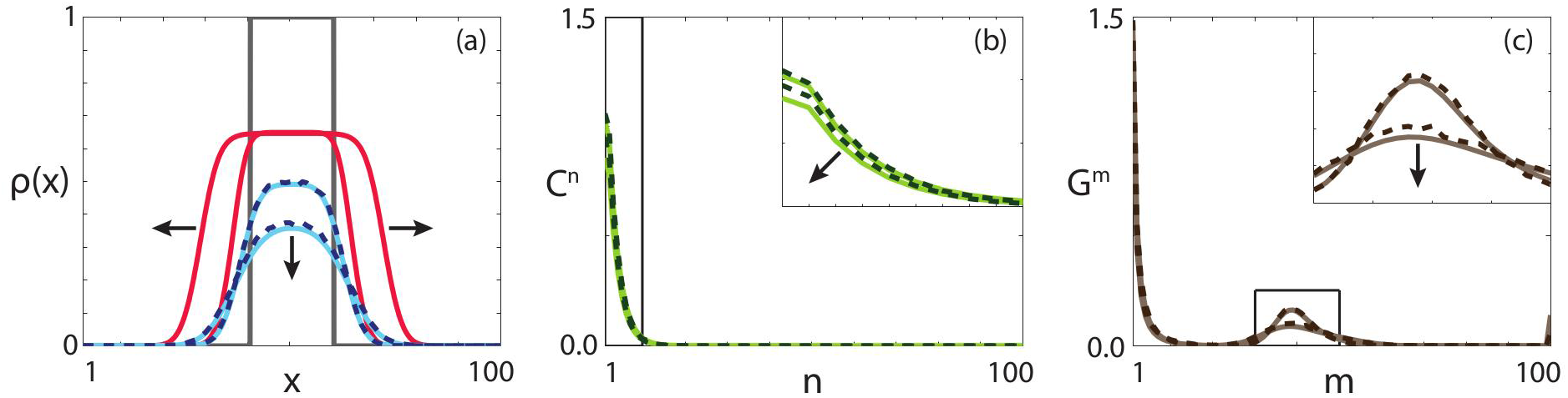
(a) Comparisons of density profiles obtained from the averaged discrete model (blue, dashed), MF description (red, solid) and C&G description (cyan, solid). (b) Comparison of the chain distribution obtained from the averaged discrete model (dark green, dashed) and C&G description (light green, solid). (c) Comparison of the gap distribution obtained from the averaged discrete model (dark brown, dashed) and C&G description (light brown, solid). All results are obtained using *P*_*m*_^1^ = 0.5, *P*_*p*_^1^= 0.45, *P*_*d*_^1^= 0.3, *P*_*m*_^*i*^= 0.25, *P*_*p*_^*i*^= 0.3, *P*_*d*_^*i*^= 0.1 for *i*∈[2, 100] *N* = 100, *M*= 10^4^, ∆ = 1. These parameter choices correspond to Special case 2. Profiles are presented at *t* = 25 and *t* = 50, and the arrow indicates the direction of increasing time. Insets highlight regions of particular interest. Grey lines represent the initial condition.

Note that all results presented here have been performed with *α* = 0, and hence no agent-agent adhesion/repulsion. Simulations performed with *α* ≠ 0 (not presented) confirm that the C&G description accurately predicts the average discrete behaviour in all cases, even with strong agent-agent adhesion/repulsion.

## Experimental case study

To highlight the insight provided about spatial clustering by the C&G description, we consider a case study motivated by a scratch assay. Scratch assays are widely used to observe the collective behaviour of a cell population in response to a model wound [42]. We present a schematic representation and experimental images of a scratch assay in Figure 6. In a scratch assay, a cell population is placed on a dish and allowed to grow to confluence. A portion of the cell population is then removed, and the remaining cells spread, through a combination of migration and proliferation, into the newly-vacant space [42]. A schematic representation of the confluent population before and after the scratch is performed is presented in Figures 6(a)-(b), with a typical experimental field of view highlighted in green. To mimic the geometry of this experiment, we consider 1 ≤ *x* ≤ 100 and initially set ρ(*x*) = 1 for *x* ≤ 30 and ρ(*x*) = 0, otherwise. Note that scratch assays are two-dimensional processes, as observed in the experimental images of a scratch assay for a 3T3 fibroblast population in Figures 6(c)-(d), and the corresponding schematics for this experiment in Figures 6(e)-(f). Full experimental details are given in [6].

**Figure 6.**
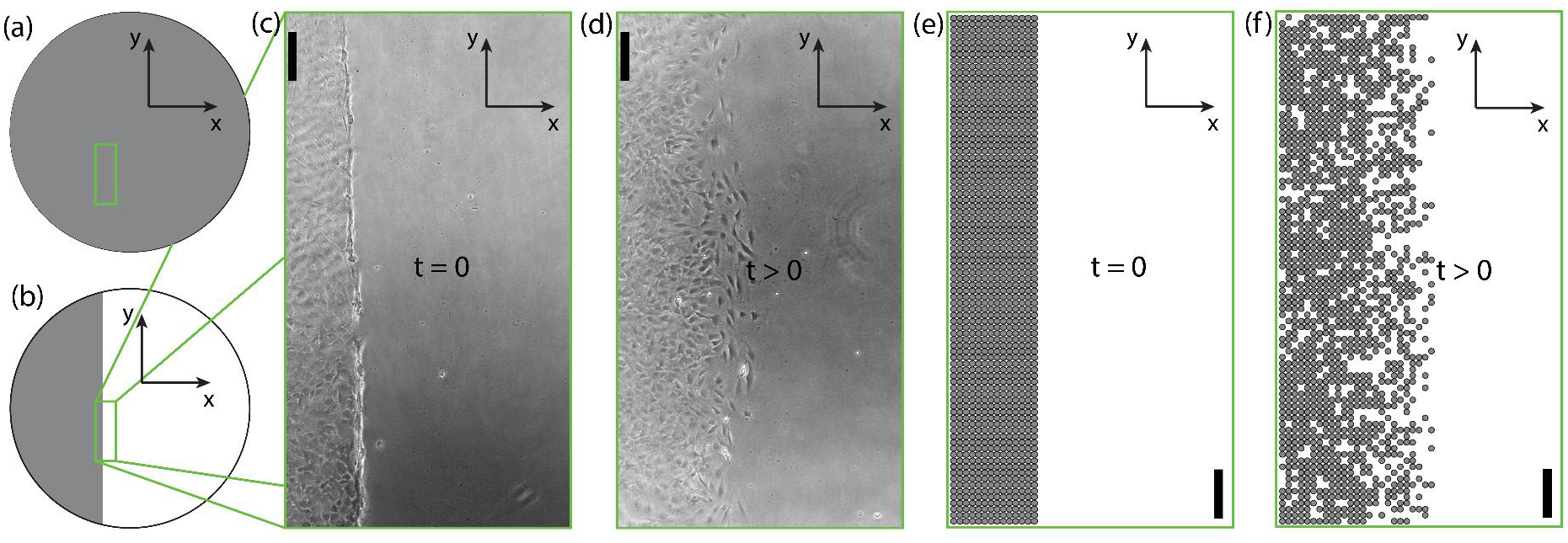
Schematic and experimental images of a scratch assay. (a) Schematic of a confluent cell population. (b) Schematic of a cell population after the scratch has been performed. Grey regions represent the confluent cell population and white regions represent the vacant area. (c)-(d) Experimental images of a 3T3 fibroblast scratch assay at (c) *t* = 0, and (d) *t >* 0. (e)-(f) Schematic of a mathematical model of a scratch assay at (e) *t* = 0, and (f) *t >* 0. Scale bar corresponds to 200 μm.

As the experiment is approximately spatially-uniform in one direction, we can approximate the scratch assay with a one-dimensional model [43]. We consider two scratch assays performed with two different cell lines where parameter estimates for the cell motility and cell proliferation rates have been presented previously: 3T3 fibroblast cells (3T3 cells) and MDA MB 231 breast cancer cells (231 cells) [44]. The investigation performed by Simpson *et al.* [44] resulted in parameter estimates of *P*_*p*_^*i*^ = 0.056 h^-1^, *P*_*m*_^*i*^ = 0.66 h^-1^ and *P*_*d*_^*i*^ = 0 h^-1^ for all *i*for 3T3 cells, and *P*_*p*_^*i*^ = 0.069 h^-1^, *P*_*m*_^*i*^ = 0.04 h^-1^, h^-1^*, P*_*d*_^*i*^=0 h^-1^ for all *i* for 231 cells. Note that the ratio 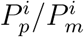 is approximately one order of magnitude higher for the breast cancer cells compared to the fibroblasts, which implies that the spatial correlations between breast cancer cells will be more significant [22].

For the numerical solution corresponding to the 3T3 cell population, presented in Figure 7, both the MF description and the C&G description approximate the average discrete behaviour reasonably well. However, the C&G description provides additional information regarding the clustering present within the migrating cell population. The chain distribution, presented in Figure 7(b), suggests that the 3T3 population does not form significant clusters, as the majority of the chains are short length.

**Figure 7.**
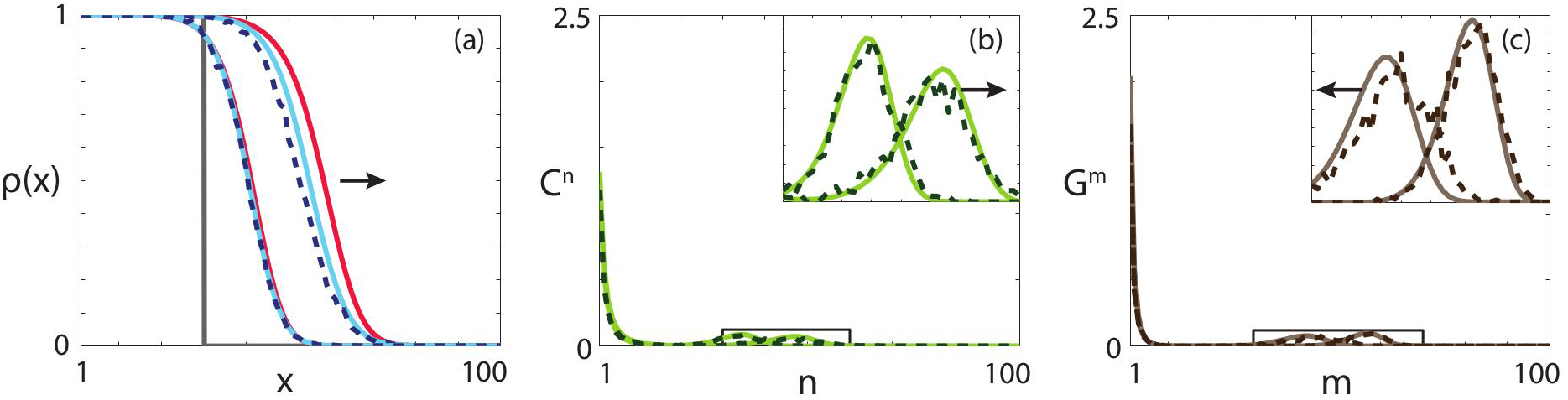
(a) Comparisons of density profiles obtained from the averaged discrete model (blue, dashed), MF description (red, solid) and C&G description (cyan, solid). (b) Comparison of the chain distribution obtained from the averaged discrete model (dark green, dashed) and C&G description (light green, solid). (c) Comparison of the gap distribution obtained from the averaged discrete model (dark brown, dashed) and C&G description (light brown, solid). All results are obtained using *P*_*m*_^*i*^ = 0.66 h^-1^, *P*_*p*_^*i*^= 0.056 h^-1^, *P*_*d*_^*i*^ = 0 h^-1^for *i* ∈ [1, 100], *N*= 100, *M* = 10^3^, ∆ = 1. Theseparameter choices correspond to 3T3 fibroblast cells. Profiles are presented at *t* = 75 and *t* = 150, and the arrow indicates the direction of increasing time. Insets highlight regions of particular interest. Grey lines represent the initial condition.

In contrast, for the numerical solution corresponding to the 231 cell population, presented in Figure 8, the C&G description accurately approximates the average discrete behaviour, while the MF description does not. This is result is intuitive as we observe that there is significantly more clustering present in the system, compared to the numerical solution corresponding to the 3T3 cell population. That is, the chain distribution in Figure 8(b) contains significantly fewer chains of short length, compared to the chain distribution in Figure 7(b). For example, at the times shown, *C*^1^ is approximately 25 times higher in the 3T3 cell population, compared to the 231 cell population. Critically, the C&G description accurately predicts the experimental observation that 231 cells form clusters while 3T3 cells do not [44]. Specifically, during monolayer formation, 3T3 cells form an approximately spatially uniform monolayer while 231 cells develop into clusters [44].

**Figure 8.**
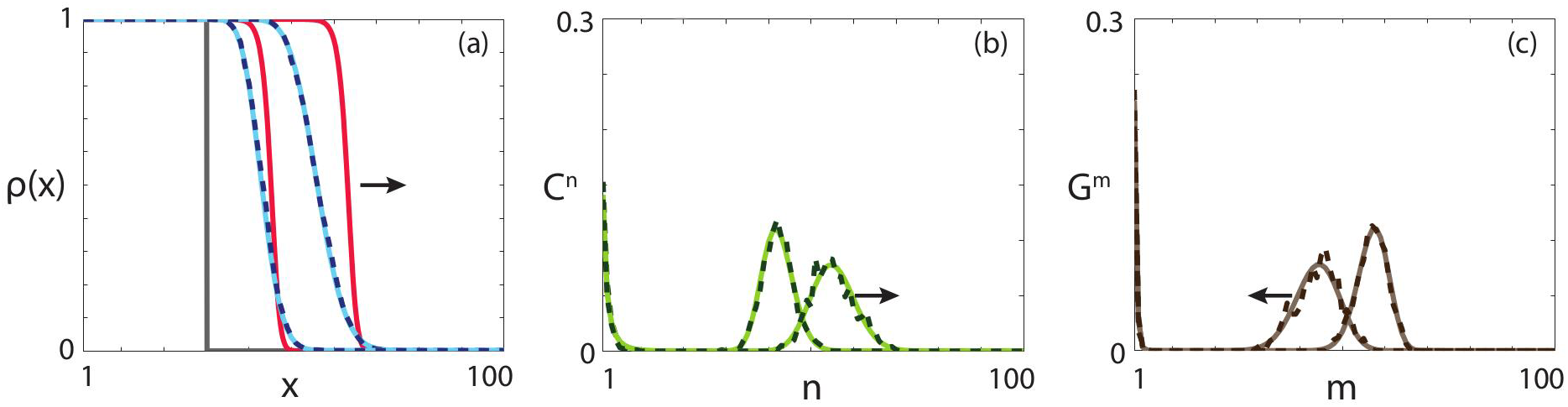
(a) Comparisons of density profiles obtained from the averaged discrete model (blue, dashed), MF description (red, solid) and C&G description (cyan, solid). (b) Comparison of the chain distribution obtained from the averaged discrete model (dark green, dashed) and C&G description (light green, solid). (c) Comparison of the gap distribution obtained from the averaged discrete model (dark brown, dashed) and C&G description (light brown, solid). All results are obtained using *P*_*m*_^*i*^ = 0.04 h^-1^, *P*_*p*_^*i*^=0.069 h^-1^, *P*_*d*_^*i*^ = 0 h^-1^ for *i* ∈ [1, 100], *N* = 100, *M* = 10^3^, ∆ = 1. These parameter choices correspond MDA MB 321 breast cancer cells. Profiles are presented at *t* = 250 and *t* = 500, and the arrow indicates the direction of increasing time. Grey lines represent the initial condition.

## Discussion and conclusions

Processes that involve moving fronts are common in cell biology and ecology [1, 2, 4, 5, 6, 7, 9, 10, 11, 12], and lattice-based random walks are widely employed to describe these processes [6, 16, 18, 19, 20, 21, 45]. Due to the stochastic nature of random walks, it can be computationally intractable to perform sufficiently many realisations of a random walk model to obtain average behaviour that is not dominated by fluctuations. Furthermore, it is difficult to extract meaningful information about population behaviour through analysis of the random walk. The standard approach to overcome these issues is to derive a MF description of the random walk [19, 20, 25]. However, this approach relies on the assumption that any spatial correlations within the random walk are weak [22, 24]. CMF descriptions that account for the spatial correlations have been proposed [22, 24, 46]. Unfortunately, these descriptions are not applicable to problems involving moving fronts as the ODEs governing the CMF description are singular in regions where the density of agents is zero [22, 24].

Here we develop and present an accurate continuum description for moving fronts associated with lattice-based random walks that contain crowding, birth, death, movement and agent-agent adhesion. We consider processes that are spatially variable, and include birth, death and movement rates that are chain-length dependent. Our C&G description provides predictions that match the average behaviour of the discrete model well in all parameter regimes. In contrast, the MF description is less flexible in terms of the birth, death and movement rates and only provides a valid approximation of the average discrete behaviour in extremely limited parameter regimes. Furthermore, for all cases considered in this work, the C&G description requires less computation time than performing 1000 realisations of the discrete model. A comparison between the time taken to perform a single realisation of the discrete model, 1000 realisations of the discrete model, and to obtain the numerical solution to the C&G system of equations is presented in Table 1.

**Table 1.**
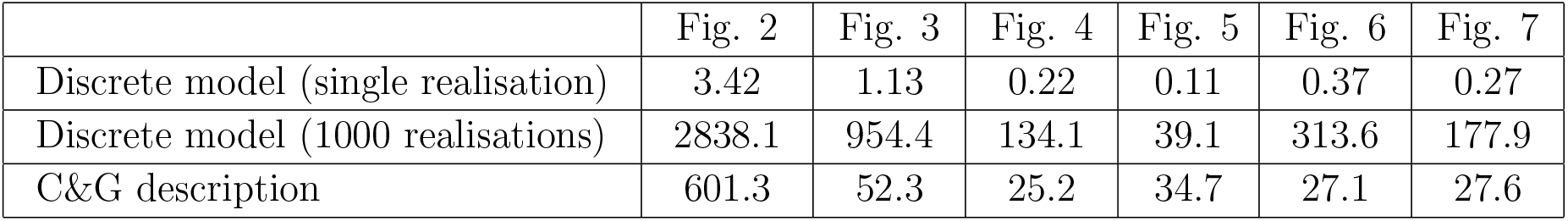
Time in seconds taken to perform: (i) a single realisation of the discrete model; (ii) 1000 realisations of the discrete model, and (iii) a numerical solution of the C&G system of equations for the parameter values in Figures 2-7. All solutions are obtained using a single 3.0 GHz Intel i7-3540M desktop processor. Note that the computation time for 1000 realisations is lower than performing 1000 repeats of a single realisation due to the time associated with initial set-up.

For the special case where the rates of birth, death and movement are independent of the chain length, the MF description correctly predicts the persistence of the population but inaccurately predicts the front velocity. For the special case where the rates of birth, death and movement are different depending on whether the agents are part of a chain of length one, or are part of a chain of length two or greater, the MF description predicts persistence when the population becomes extinct, and predicts extinction when the population persists. In both these cases the C&G description accurately predicts the front velocity, and the persistence or extinction of the population. Furthermore, the C&G description provides information about the spatial clustering of both occupied and unoccupied sites, and the clustering predictions approximate the clustering observed in the discrete model accurately.

The work presented here could be extended in several ways. The influence of local spatial structure on the persistence of species is a key question in ecology [34, 35, 45, 47]. The C&G description provides an explicit estimate of the spatial clustering of both agents and unoccupied space. Therefore, it would be instructive to apply the C&G description to ecological processes to obtain insight into the clustering present in the system for parameter regimes where the agent population becomes extinct. Another approach would be to investigate a truncated system of governing equations, where there is a maximum chain or gap length. If there is prior knowledge about the long-time density of the system, an assumption could be made that chains or gaps above a threshold length could be neglected, and hence the system of equations could be truncated. This truncation would reduce the computational cost associated with obtaining a numerical solution to the governing equations. It would be insightful to examine the trade-off between the reduction in computational cost and the decreased accuracy caused by the truncation. Alternatively, the C&G description presented here could be calibrated to experimental data from the cell biology literature. For example, lattice-based random walks have been calibrated to *in vitro* cell biology experiments to obtain estimates of cell diffusivity and cell proliferation rates [6, 21]. However, the calibration of stochastic models to experimental data is computationally expensive [6, 48]. As the C&G description accurately approximates the average random walk behaviour in all parameter regimes, it would be instructive to determine whether similar cell diffusivity and cell proliferation rates could be obtained from calibration of the deterministic C&G description to experimental data, and to quantify the reduction in computation time to obtain the parameter estimates. However, we leave these extensions for future consideration.

## Acknowledgements

We appreciate the assistance provided by Emeritus Professor Sean McElwain. This work is supported by the Australian Research Council (FT130100148, DP140100249). We thank two referees and the editor for their helpful comments.

## Appendix A. Governing equations

### Appendix A.1. Chains

Here we present the time rate of change for each chain length and location, obtained by considering the potential outcomes of each type of birth, death and movement event.

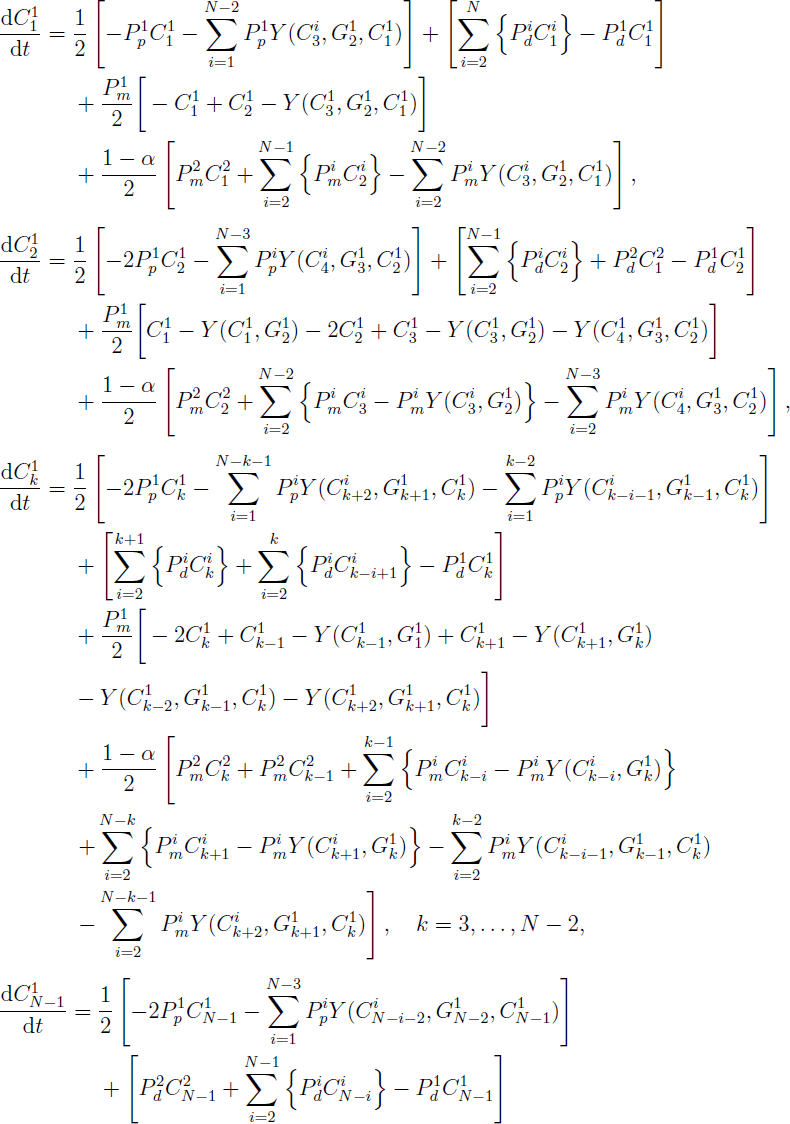

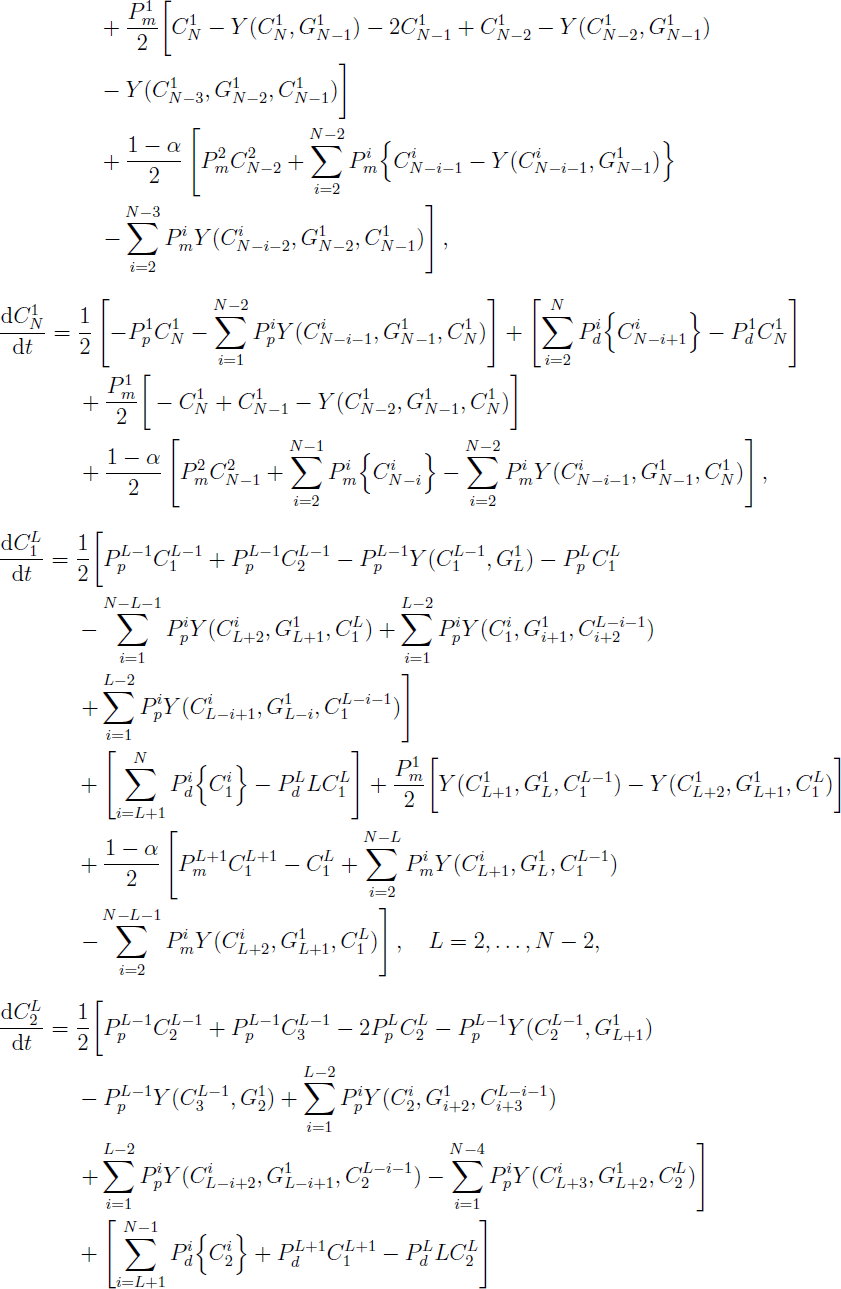

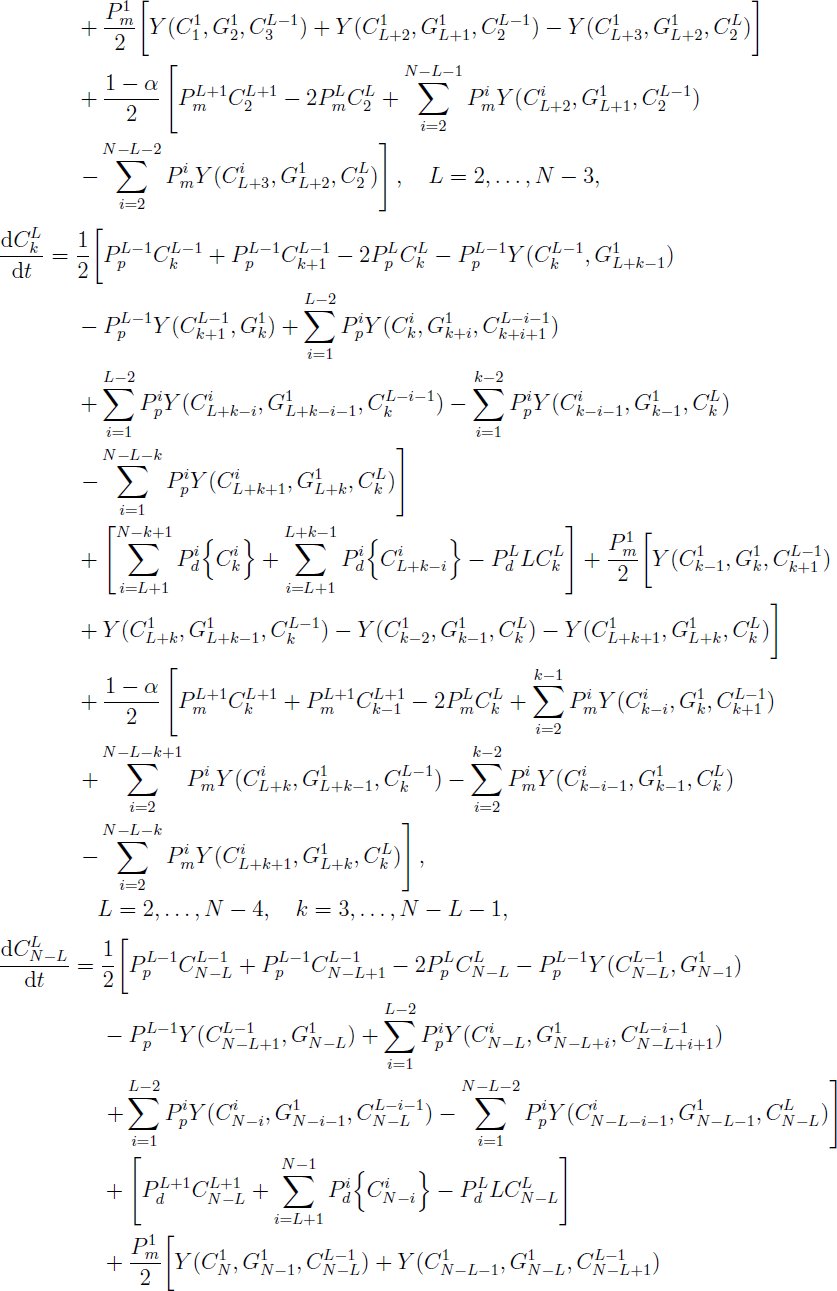

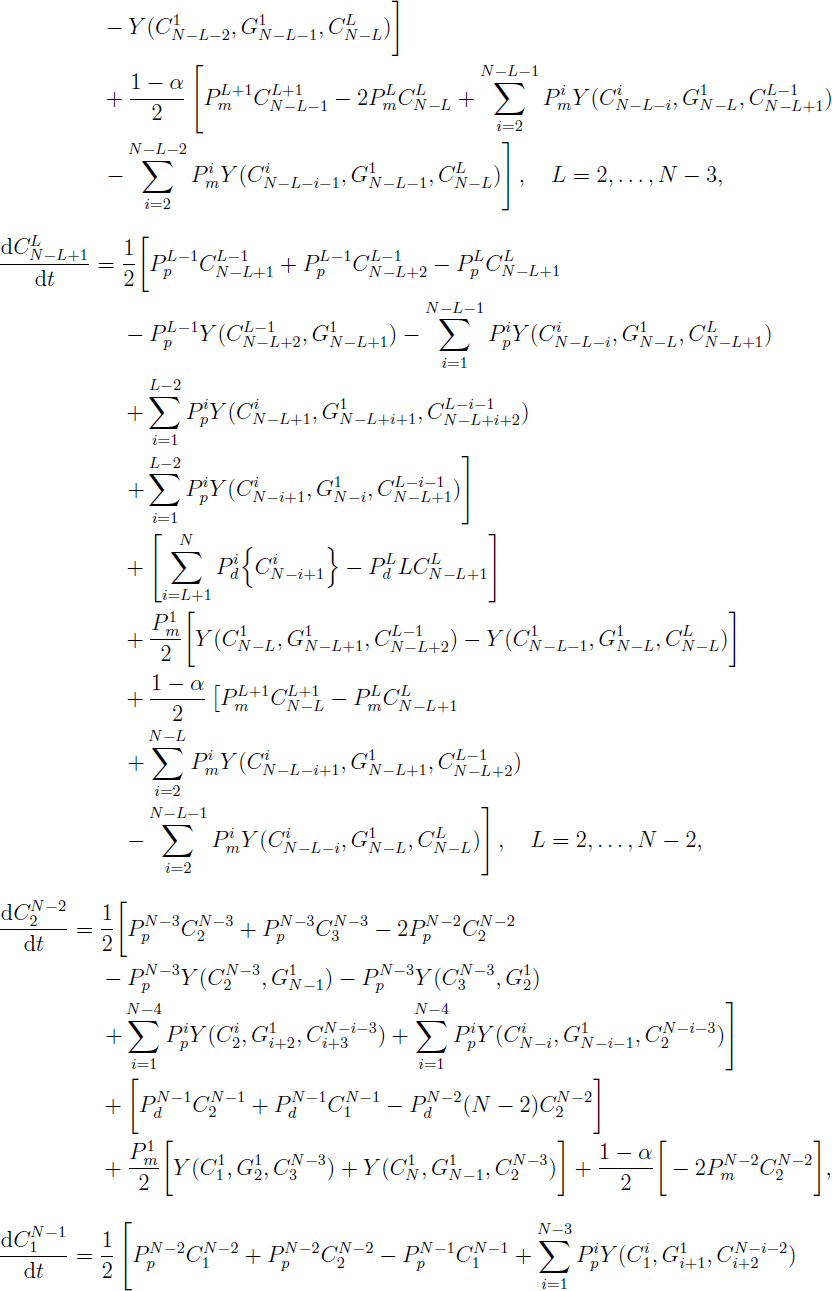

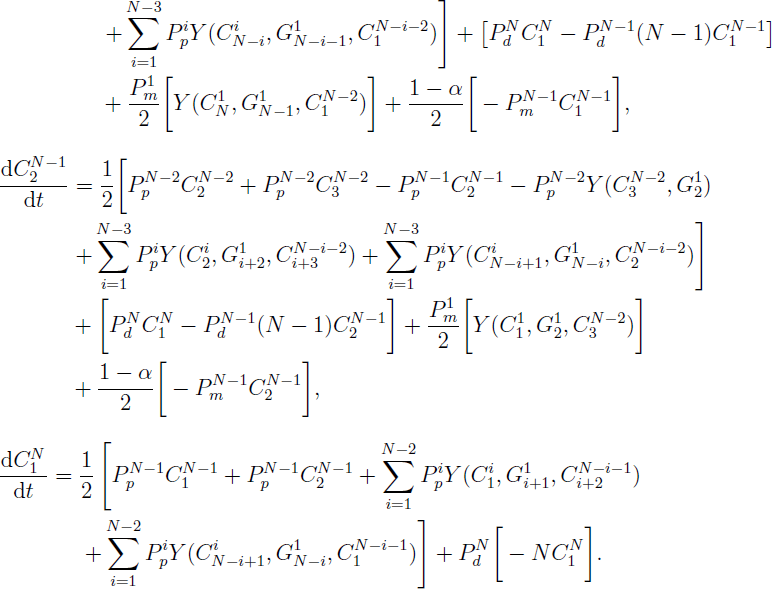

### Appendix A.2. Gaps

Here we present the time rate of change for each gap length and location, obtained by considering the potential outcomes of each type of birth, death and movement event.

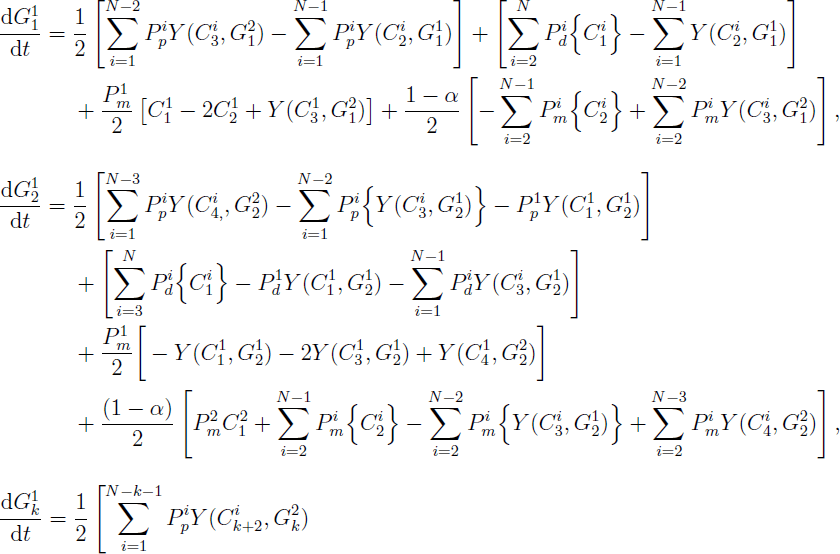

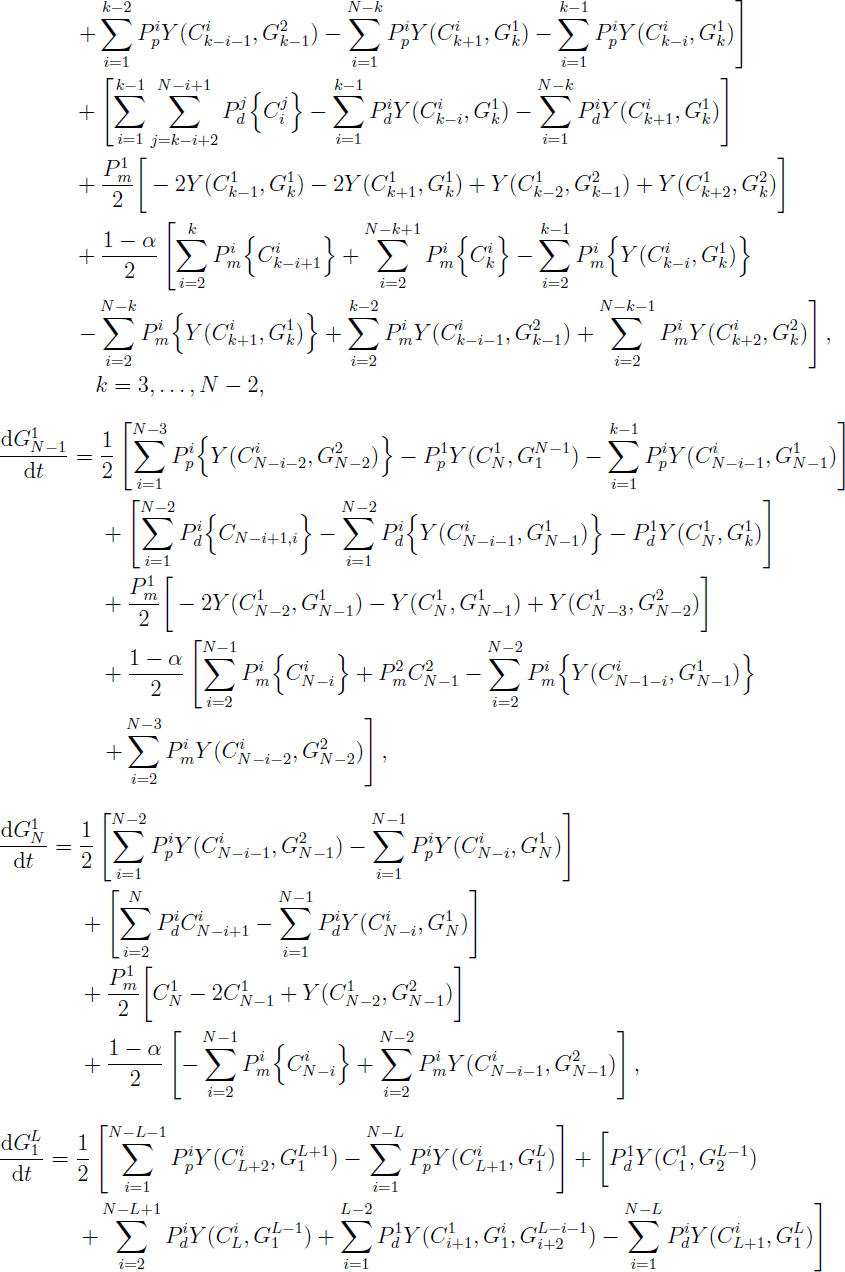

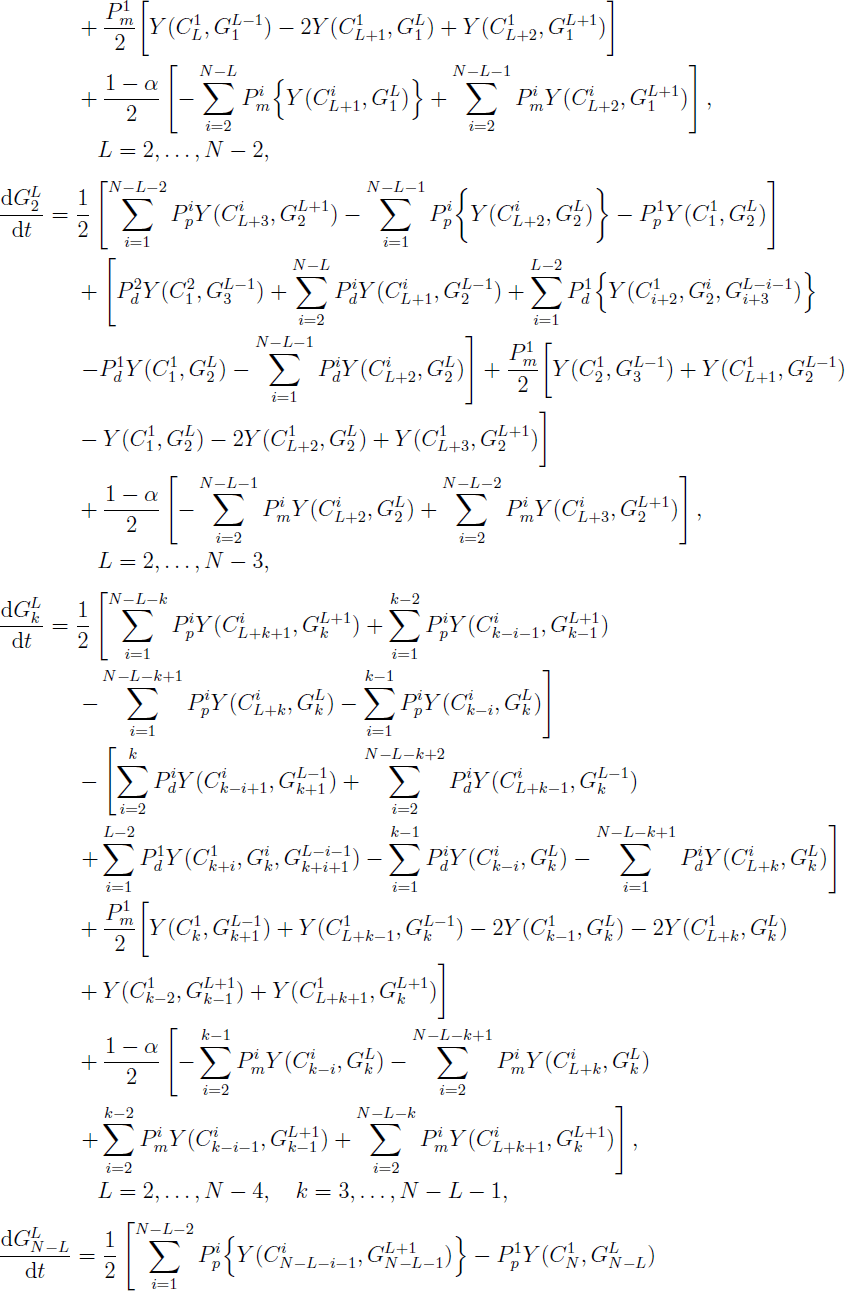

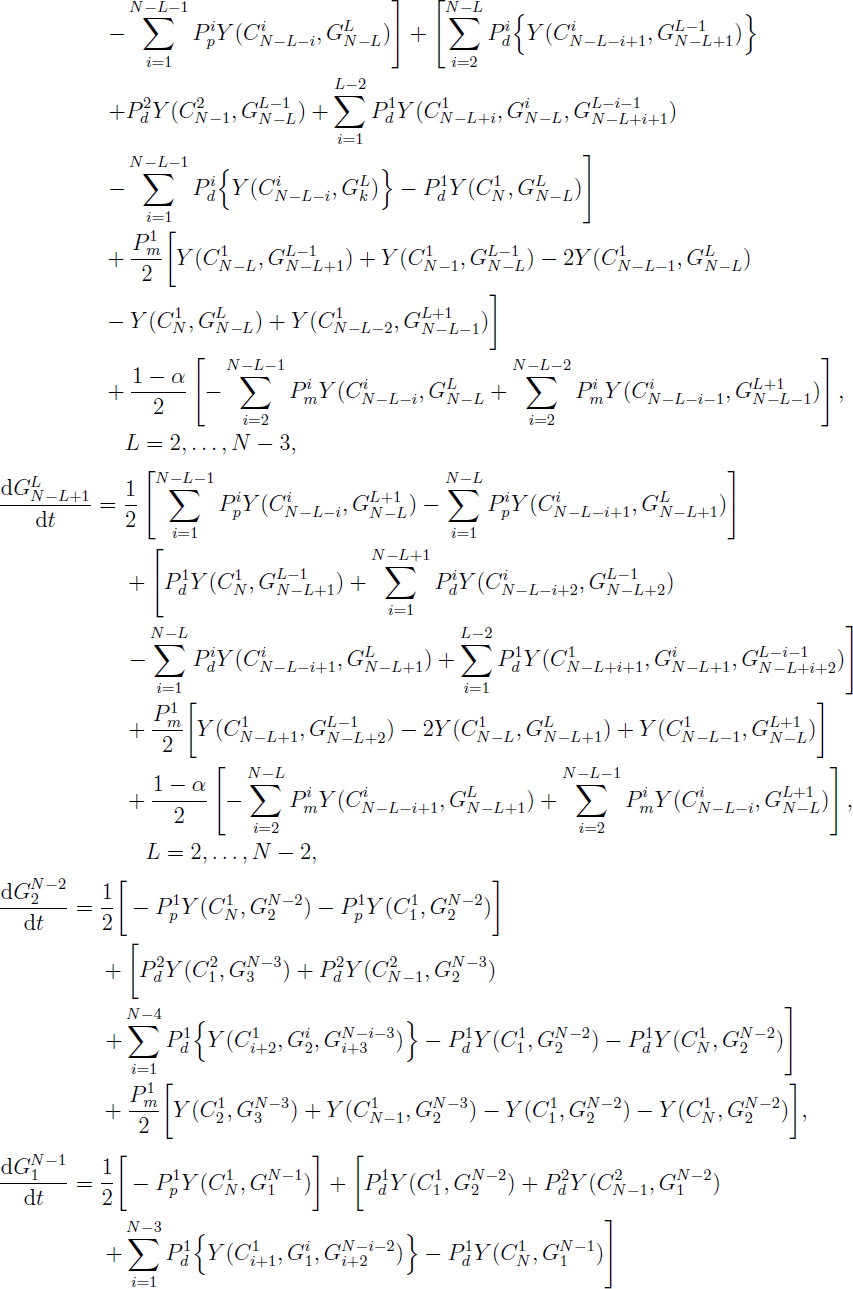

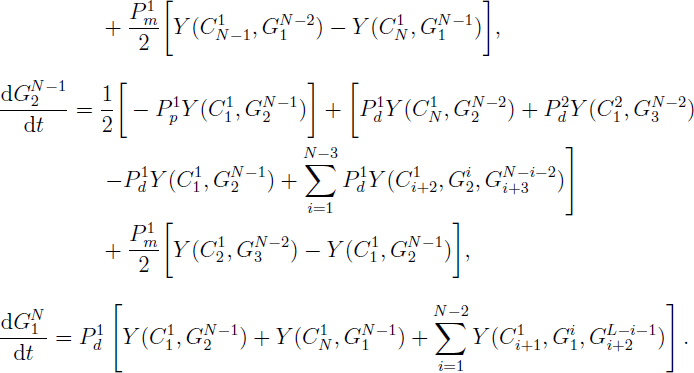

The *Y* function represents the number of configurations that contain the specified chains and gaps within the parentheses, and is defined as

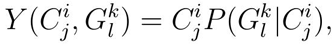

where 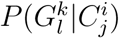 represents the probability that a gap of length *k* at site *l* exists, given that there is a chain of length *i* at site *j*, and can be calculated as

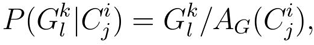

where *A*_*G*_(*C*_*j*_^*i*^) are all the possible gaps that can exist on either side of the chain of length *i* at site *j*. If we are interested in gaps on the positive *x*side of the chain, we require *l*= *i* + *j*and *k ≤ N − i − j*+1. If we are interested in gaps on the negative *x* side of the chain, we require *l≤j*-1, *k* = *j-k*. Similarly,

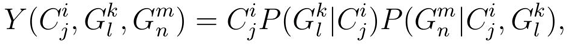

where 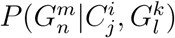 is the probability a gap of length *m* at site *n* exists, given that there is a chain of length *i* at site *j* and a gap of length *k* at site *l*. Note that the gaps at site *l* and site *n* must necessarily be on opposing sides of the chain at site *j* and hence 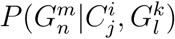 is calculated in the same manner as 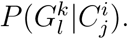.

## Appendix B. Numerical techniques

### Appendix B.1. Ordinary differential equations

The system of ODEs describing the dynamics of the chains and gaps, presented in Appendices A.1 and A.2, are solved using an adaptive Runge-Kutta method with a strict truncation error control of 10^-6^ [49]. All results presented are found to be insensitive to a reduction in the strict truncation error control.

### Appendix B.2. Partial differential equations

The two mean-field descriptions of the discrete process, Equation (1) and Equation (2), are discretised onto a spatially uniform finite difference grid with grid spacing *δx*. The spatial derivative terms are approximated using a central finite difference approximation. We approximate the temporal derivative using the backward Euler method with constant time step *δt*, and the resulting system of nonlinear algebraic equations is solved using Picard iteration with absolute convergence tolerance *ϵ*. Finally, the system of tridiagonal algebraic equations is solved using the Thomas algorithm [49]. In all cases, δ*x* = 0.1, δ*t* = 0.01 and ϵ = 10^−6^.

